# Single-cell analysis identifies EpCAM^+^/CDH6^+^/TROP-2^−^ cells as human liver progenitors

**DOI:** 10.1101/294272

**Authors:** Joe M Segal, Daniel J Wesche, Maria Paola Serra, Bénédicte Oulès, Deniz Kent, Soon Seng Ng, Gozde Kar, Guy Emerton, Samuel J I Blackford, Spyros Darmanis, Rosa Miquel, Tu Vinh, Ryo Yamamoto, Andrew Bonham, Alessandra Vigilante, Sarah Teichmann, Stephen R. Quake, Hiromitsu Nakauchi, S Tamir Rashid

## Abstract

The liver is largely composed of hepatocytes and bile duct epithelial cells (BECs). Controversy exists as to whether a liver stem/progenitor cell capable of renewing both hepatocytes and BECs exists. Single cell RNA sequencing of freshly isolated human foetal and healthy adult liver identified hepatocyte, hepatoblast and liver progenitor cell (hLPC) populations. hLPCs, found at the interface between hepatocytes and bile ducts in both foetal and adult tissue, were distinguishable from BECs by their negative expression of TROP-2. Prospective isolation followed by in vitro culture demonstrated their potential for expansion and bi-lineage differentiation. The hLPC expression signature was also conserved within expanded cell populations specific to certain cases of liver injury and cancer. These data support the idea of a true progenitor existing within healthy adult liver that can be activated upon injury. Further work to define the mechanisms regulating hLPC behaviour could advance understanding of human development and disease.

## Introduction

The liver has a remarkable capacity to regenerate. Conflicting evidence exists regarding the cellular origin of this process and mechanisms at play. Contradictory findings may in part be attributed to differences between regeneration seen at times of normal homeostasis versus times of injury. In rodent models for example the regenerative potential has been attributed to hepatocytes (Yanger et al., 2014, Schaub et al., 2014), cholangiocytes (Tarlow et al., 2014), stem/progenitor cells located in the ductal region (Lu et al., 2015, Raven et al., 2017, Petersen et al., 1998, Huch et al., 2013, Furuyama et al., 2011)(Dabeva et al., 2000) (Yovchev et al., 2008), as well as stem/progenitor cells located around the central vein (Wang et al., 2015). In humans it has been proposed that hLPCs derived from EpCAM^+^ cells found in the ductal plate during foetal liver development, localise to the canals of Hering after birth (Schmelzer et al., 2007) and become reactivated at times of severe chronic liver injury forming what is pathologically described as ductular reactions (Lowes et al., 1999). Recently it was proposed that EpCAM^+^ liver cells, like stem cells in other adult tissues, are LGR5^+^ WNT responsive cells that can be isolated from human adult liver, serially expanded *ex vivo* and still retain their potential for differentiation (Huch et al., 2013). Given that EpCAM is a cell surface marker likely common to multiple cell types we hypothesized that previously labelled EpCAM^+^ hLPC represented a heterogeneous population of cells. Single cell RNA sequencing (scRNA-seq) is an unbiased tool to characterise the transcriptome of a single cell which has already been used to uncover the presence and function of novel cell types, and the existence of transient cell states, within previously undefined heterogeneous populations (Treutlein et al., 2014, Darmanis et al., 2017, Philippeos et al., 2018, Proserpio et al., 2016). We therefore employed this tool to interrogate the cellular heterogeneity within EpCAM^+^ populations of human liver tissue to identify a true liver stem / progenitor cell.

## Results

### Enrichment for progenitor cells in human foetal liver

Previous studies identified putative human hepatic stem /progenitor cells to be within the EpCAM^+^ /NCAM^+^ population (Schmelzer et al., 2007). We therefore decided to employ a cell sorting strategy to enrich for potential hLPCs and deplete contaminating blood cells. We obtained five separate freshly isolated human foetal liver samples during the second trimester of gestation. Cells were isolated as previously reported (Gramignoli et al., 2012) and sorted on cell surface markers CD235a (red blood cell marker), CD45 (leukocyte marker), EpCAM and NCAM. Only CD235a^−^/CD45^−^; CD235a^−^/CD45^−^/EpCAM^+^/NCAM^−^ and CD235a^−^/CD45^−^/EpCAM^+^/NCAM^+^ (Figure 1A, Supplementary Figure 1Ai) were used for downstream analysis (Supplementary table 1). RNA from each cell was isolated and sequenced across four 384-multiplex libraries (along with adult single cell samples). In total (including adult cells, Figure 3A) ~1400 cells were captured and sequenced. Post-alignment, raw counts were processed through the SCATER scRNA pipeline for quality control (qc) (McCarthy et al., 2017). After SCATER analysis 384 foetal cells passed qc based on the median absolute deviation (MAD) of genes expressed (%), library size (millions) (Supplementary Figure 1Bi), mitochondrial gene expression (%) and artificial ERCC spike-in gene proportion (%) (Supplementary Figure 1Ci). Sample counts were normalised by transcripts per million (TPM).

**Figure 1:**
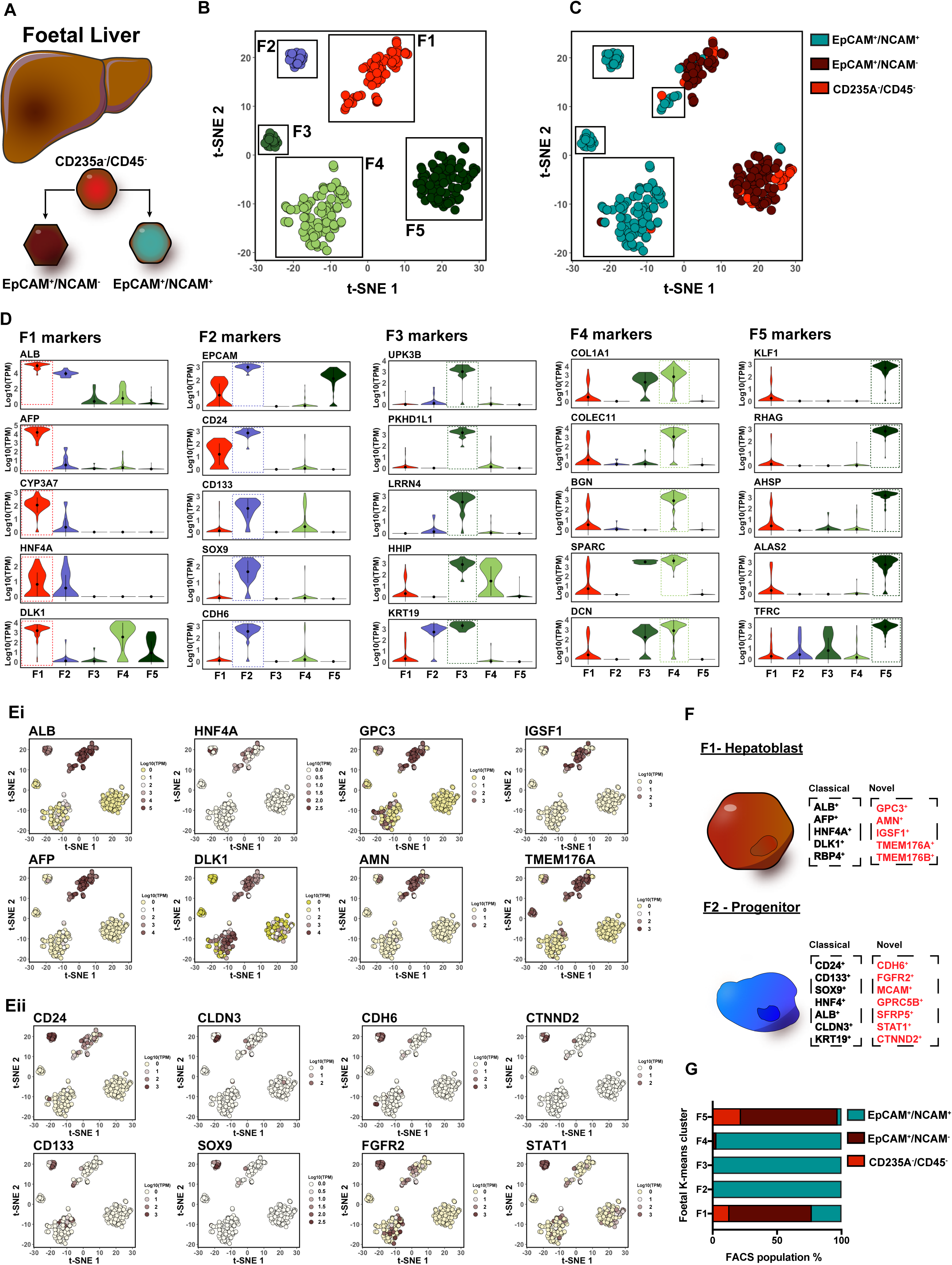
ScRNA-Seq of foetal liver EpCAM^+^/NCAM^+^ cells identifies a sub-population of human liver progenitor cells in foetal liver. (A) Overview of FACS strategy in foetal human liver to collect cells for scRNA-seq. (B) K-means clustering applied to 2D t-SNE visualisation of single cells isolated from foetal human liver. T-SNE plot points are differentially coloured by cluster and labelled as F1-F5. (C) 2D t-SNE visualisation of single cells isolated from foetal human liver included in the study. T-SNE plot points coloured by FACS parent population. (D) Violin plots detailing expression levels of selected marker genes in each cluster identified in foetal liver scRNA-seq analysis. Expression is Log10(TPM). (E) Expression of selected enriched F1 (i) and F2 (ii) marker transcripts overlaid on the 2D t-SNE space of populations identified in foetal liver scRNA-seq analysis. (F) Schematic of known and novel markers expressed in foetal clusters F1 and F2 labelled as ‘Hepatoblast’ and ‘Progenitor’ cells respectively. (G) Bar-plots representing the percentage abundance of each FACS populations for the foetal populations F1-F5.

### Unbiased transcriptome-wide gene expression profiling identifies a putative human liver progenitor within EpCAM^+^/NCAM^+^ double positive FAC-sorted population

After sequencing, we employed t-Distributed Stochastic Neighbour Embedding (t-SNE) on high variance genes, combined with K-means clustering analysis, identifying five total clusters (labelled F1-F5) across all FACS sorted foetal liver cells (Figure 1B). EpCAM^+^/NCAM^+^ cells were identified in four of the five foetal populations (Figure 1C). Only clusters F1 and F2 were enriched for ALB expression, defining them to be hepatic (Doweiko and Nompleggi, 1991) (Figure 1D). The remaining clusters, F3, F4 and F5 were negative for *ALB* and *HNF4a* suggesting the presence of non-hepatic cell types (Ballester et al., 2014). Hepatoblast markers *AFP* and *DLK1* (Tanaka et al., 2009, Kuhlmann and Peschke, 2006) were enriched in population F1. Population F2 expressed previously reported hepatic progenitor markers *EpCAM*, *CD24*, *SOX9* and *CD133* (Furuyama et al., 2011, Yovchev et al., 2008), but were negative for AFP. Population F3 expressed markers of mesothelial cells such as *UPK3B* and *PKHD1L1* (Kanamori-Katayama et al., 2011). Both populations F2 and F3 were enriched for the LPC marker KRT19 (Tarlow et al., 2014). F3 also expressed the mesenchyme marker *COL1A1* (Friedman, 1996). F4 expressed *COL1A1* alongside other extracellular matrix (ECM) associated genes including *DCN* and *BGN* (Rashid et al., 2012). F5 expressed exclusive markers of erythroid cells such as *KLF1* and *RHAG* (Miller and Bieker, 1993) (Hu et al., 2013) (Figure 1D). This was unexpected given our rigorous FACS strategy to deplete erythroid cells, informed by previous literature (Schmelzer et al., 2006). We therefore confirmed through flow cytometric analysis that the F5-specific marker *TFRC* was expressed on a sub-population of CD235a^−^ and EpCAM^+^ cells (Supplementary Figure 1D), confirming that the F5 population was not a contaminate cell-type. This suggests the possibility of erythroid-like cells existing within CD235a^−^/EpCAM^+^ foetal liver tissue.

Alongside the classical hepatoblast markers, F1 cells were also enriched for unreported human markers *GPC3*, *AMN*, *IGSF1* and *TMEM176A* (Figure 1Ei, Figure 1F, Supplementary Table 2). F2 was similarly enriched for potential new hepatic progenitor markers: *CDH6*, *CTNND2*, *FGFR2* and *STAT1* (Figure 1Eii, Figure F, Supplementary Table 3). Interestingly, the F2 (putative progenitor) population was derived exclusively from double positive EpCAM^+^/NCAM^+^ cells, while F1 (hepatoblasts) came predominantly from the EpCAM^+^/NCAM^−^ (EpCAM only) population. Cells in F3 and F4 populations were also derived from the EpCAM^+^/NCAM^+^ double positive population, suggesting significant heterogeneity exists within cell populations previously identified to be hepatic stem/progenitor cells (Huch et al., 2015, Schmelzer et al., 2007) (Figure 1G).

### CDH6 defines a sub population of EpCAM^+^/NCAM^+^ cells as true progenitors

Having computationally established the presence of a novel CDH6^+^ progenitor cell type, we went on to investigate its tissue specific localisation in foetal liver. Using *in situ* RNA hybridisation (RNA-ISH), we identified *CDH6* and other defining markers *FGFR2*, *CTNND2* and *STAT1* enriched in the ductal plate (Terada, 2013), a layer of cells surrounding the portal tract (Figure 2A, Supplementary Figure 2A). Protein expression was validated by immunohistochemistry (IHC) and immunofluorescence (IF). We confirmed these regions were positive for previously reported ductal plate and hepatic progenitor markers EpCAM and CK19 and CLDN3 (Terada, 2013, Schmelzer et al., 2006), along with our newly identified markers CDH6 and STAT1 (Figure 2B-C, Supplementary Figure 2B).

**Figure 2:**
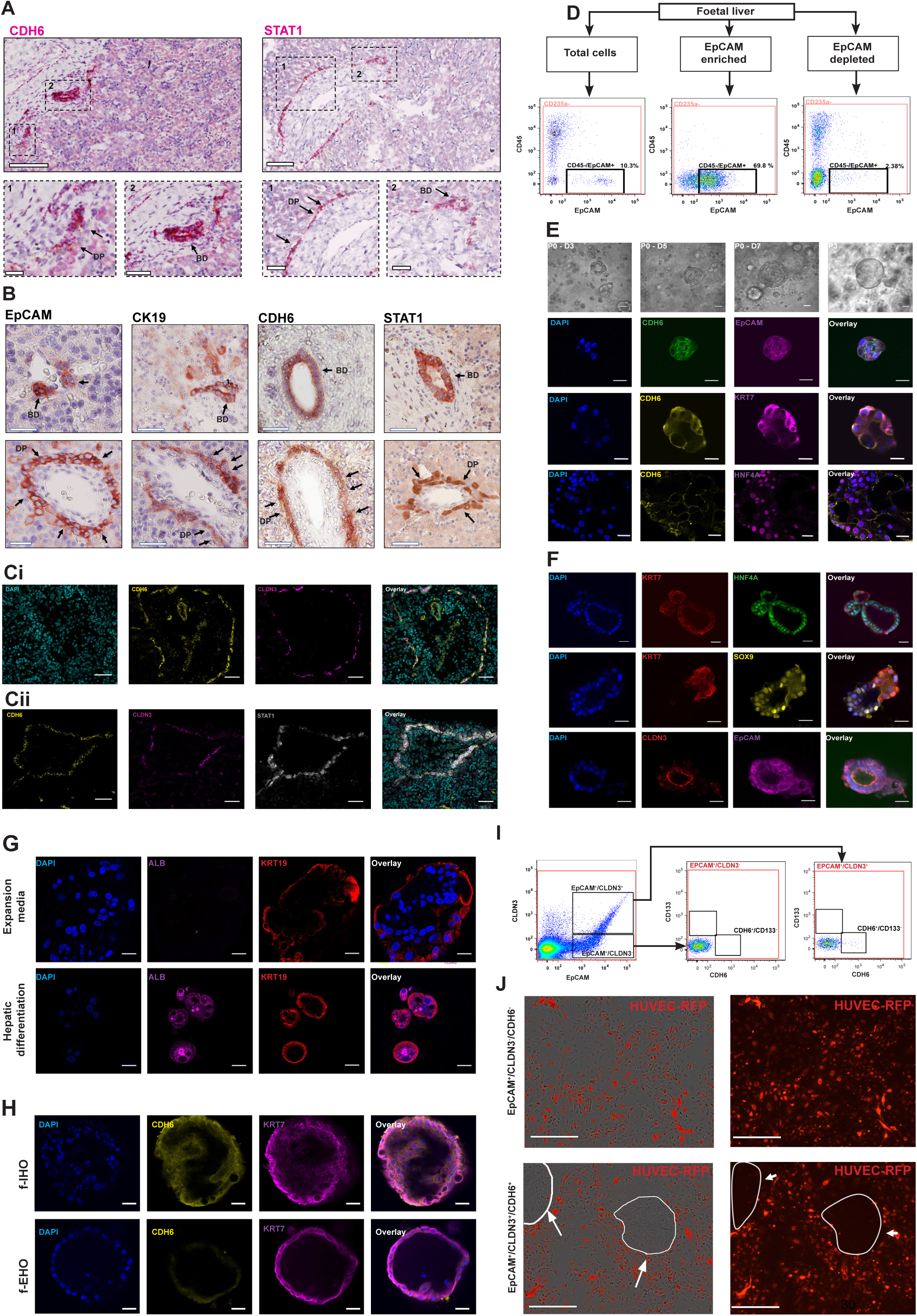
CDH6^+^ human liver progenitor cells reside in the ductal plate of foetal liver. (A) RNA-ISH for CDH6 and STAT1 on human foetal liver ductal plate (DP) and bile ducts (BD) regions (detail of the red dots expanded in the squares). Scale bars represent 50 μm, 25 μm in blown up squares. (B) Immunohistochemistry (IHC) of EpCAM, CK19, CDH6 and STAT1 in BD and DP regions of human foetal liver. Scale bars represent 50 μm. (C) Immunofluorescence (IF) staining of CDH6 (yellow) and CLDN3 (magenta) (i) and CDH6 (yellow), CLDN3 (magenta) and STAT1 (grey) (ii) co-expression in human 2^nd^ trimester (15-21 pcw) foetal liver slides. Slides counterstained in DAPI (cyan). (D) Representative FACS plot of MAC-sorted total (left), EpCAM enriched(middle) and EpCAM depleted (right) cells from dissociated human foetal liver. All gates were set on fluorescence minus one (FMO) controls Gate percentages are representative of n = 3 isolations. (E) Phase-contrast and IF staining of foetal intra-hepatic organoids (f-IHOs) derived from EpCAM enriched foetal liver cells in liver expansion (LE) media. All structures are counterstained with DAPI (blue). P0 refers to cells isolated directly from primary tissue. All IF staining performed between passage 3-5. (F) IF staining of f-IHOs in LE media. All structures are counterstained with DAPI (blue). All IF staining performed between passage 3-5. (G) IF staining of f-IHOs for albumin (magenta) and KRT19 (red) cultured for 7 days after passage in LE media or Hepatic differentiation (HD) media. (H) IF staining of f-IHOs and foetal ex-hepatic organoids (f-EHOs) derived from foetal gallbladder for CDH6 (yellow) and biliary marker KRT7 (magenta). Counterstained with DAPI (blue). (I) Representative scatter plot of FAC-sorting strategy for isolation of EpCAM^+^/CLDN3^+^/CDH6^+^ cells. Top plot displays EPCAM (X-axis) vs CLDN3 (y-axis) of Live/CD235a^−^/CD45^−^ negatively selected cells. Middle plot displays CDH6 (x-axis) and CD133 (y-axis) of EPCAM^+^/CLDN3^+^ cells. Bottom plot displays CDH6 (x-axis) and CD133 (y-axis) of EPCAM^+^/CLDN3^−^ cells. All plots gated on fluorescence minus one (FMO) staining controls. (J) Phase contrast and IF imaging of FAC-isolated EpCAM^+^/CLDN3^−^/CDH6^−^/CD133-(i) (scale bars represent 300 μm) and EpCAM^+^/CLDN3^+^/CDH6^+^/CD133^−^ (ii) (scale bars represent 200 μm) cells cultured for 10 days on RFP-HUVECs in LE media, with corresponding IF images. RFP negative regions circled in white. All scale bars represent 25 μm unless stated otherwise. FACS plots are representative of at least 3 independent experiments.

To validate the presence of CDH6^+^ progenitor cells in *ex vivo* culture expanded tissue, we prospectively enriched EpCAM^+^ cells from fresh foetal liver cultured in modified organoid forming conditions (Huch et al., 2015) (Figure 2D). After multiple passages, expanded organoids were positive for CDH6 as well as classical progenitor markers EpCAM, SOX9, CLDN3, KRT7 and HNF4A (Figure 2E-F). Transferring organoids to a hepatic differentiation medium, could induce expression of albumin protein after 7 days, confirming the hepatic differentiation potential of CDH6^+^ organoids (Figure 2G). In comparison, organoids derived from foetal gall bladder were negative for CDH6, positive for biliary marker KRT7 (Figure 2H), and unable to generate albumin positive cells in modified hepatic differentiation conditions (Huch et al., 2013). Taken together these results suggested that a CDH6^+^ subpopulation of foetal EpCAM^+^ cells are in fact the true hLPCs. To confirm this, we compared the expansion capability of CDH6^+^ vs CDH6-cells. EpCAM^+^/CLDN3^+^/CDH6^+^ and EpCAM^+^/CLDN3^−^/CDH6^−^ sorted cells from freshly isolated human foetal liver were seeded on RFP-labelled human umbilical vein endothelial cells (HUVEC) in liver expansion media (Figure 2I). Only CDH6^+^ cells were capable of expansion and passage off supporting HUVECS, whilst retaining EpCAM and CDH6 expression (Figure 2J, Supplementary Figure 3).

**Figure 3:**
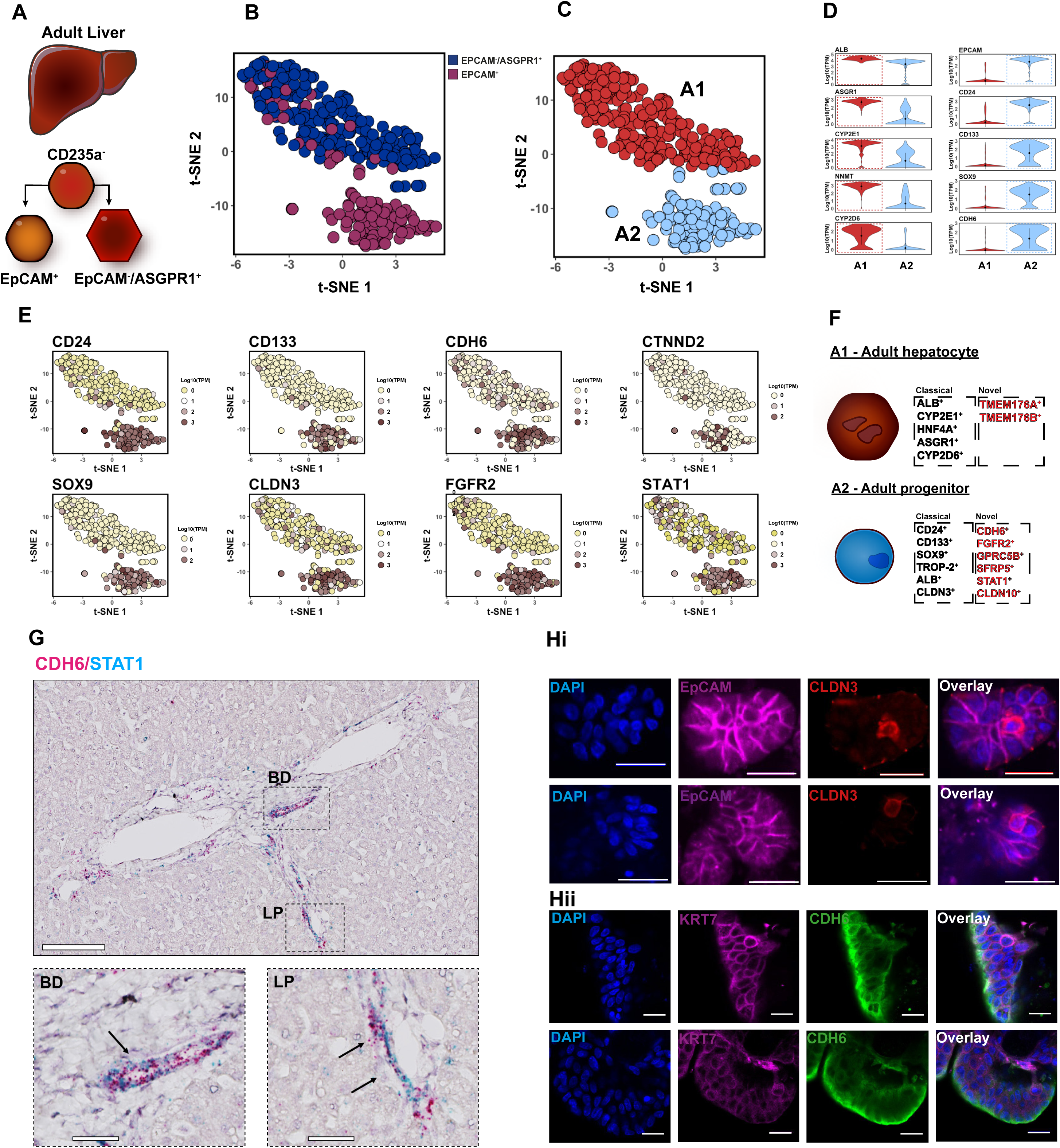
ScRNA-Seq of FAC-sorted normal adult human liver identifies a conserved progenitor within the limiting plate. (A) Overview of FACS strategy in adult human liver to isolate cells for scRNA-seq. (B) 2D t-SNE visualisation of single cells isolated from adult human liver included in the study coloured by FACS parent population. (C) 2D t-SNE visualisation of single cells isolated from adult human liver t-SNE plot points coloured by K-mean cluster and labelled A1-A2. (D) Violin plots detailing expression levels of selected marker genes in each cluster identified in adult liver scRNA-seq analysis detailing distinct cell types. Scale is log10 transcripts per million (TPM). (E) Expression of selected enriched ‘Adult progenitor’ (A2) marker transcripts overlaid on the 2D t-SNE space of adult liver scRNA-seq analysis. (F) Schematic of known and novel markers expressed in adult clusters A1 and A2 labelled as ‘Adult hepatocyte and ‘Adult progenitor’ cell types respectively. (G) Duplex RNA-ISH for CDH6(red)/STAT1(blue) in adult human intra-hepatic bile ducts (BD) and limiting plate (LP), detail of the red/blue dots expanded in the square. Scale bars represent 50 μm, 25 μm in blown up squares. (I) Immunofluorescence (IF) staining on Endoscopic retrograde cholangiopancreatography procedure (ERCP) intra-hepatic biliary brushing primary human samples for EpCAM (purple) with CLDN3 (red) (i) and KRT7 (magenta) with CDH6 (green) (ii), after 2 days in Matrigel with liver expansion (LE) media. Scale bars represent 25 μm.

### Single cell analysis of normal adult human liver identifies CDH6^+^ cells at the limiting plate

To characterise hepatic cell type diversity within adult human liver we isolated EpCAM^+^ and EpCAM^−^/ASGPR1^+^ cells obtained from fresh tissue. ASGPR1 is a marker of mature hepatocytes (Schwartz et al., 1981). This tissue was obtained from donor grafts, the remainder of which were successfully implanted into patients with liver failure (Figure 3A, Supplementary Figure 1Aii). In total 480 cells were isolated from 3 separate adult liver samples, with 357 cells passing qc (Supplementary table 1, Supplementary figure 1Bii & Cii). Analysis of transcript expression of adult scRNA-seq populations revealed a mature hepatocyte population (A1) predominantly isolated from EpCAM^−^/ASGPR1^+^ cells (Figure 3 B-C). This population highly expressed cytochrome P450 enzymes *CYP2E1*, *CYP2D6*, and the metabolic enzyme *NNMT* (Baxter et al., 2015, Aksoy et al., 1994)(Figure 3D, Supplementary Table 4). Population A2 cells, almost exclusively EpCAM^+^, were enriched for known progenitor markers *CD24*, *CD133* and *SOX9* (Figures 3D). Similarly, to the foetal hLPC population, adult hLPCs also expressed *CDH6*, *CTNND2*, *FGFR2* and *STAT1* (Figure 3E-F Supplementary Figure 4, Supplementary Table 5). RNA-ISH confirmed *CDH6* and *STAT1* expression in both bile ducts and at the limiting plate, a layer of hepatic cells that borders the portal tract (Turner et al., 2011, Paku et al., 2005) (Figure 3G). As we were unable to source further normal human adult liver tissues for prospective isolation of the CDH6^+^ population, we validated their presence in cells expanded from Endoscopic Retrograde Cholangio-Pancreatography (ERCP) brushings (Ferrari Júnior et al., 1994). We confirmed CDH6 co-expression with the classical ductal / progenitor markers CLDN3, EpCAM and KRT7 in these primary adult samples (Figure 3H).

**Figure 4:**
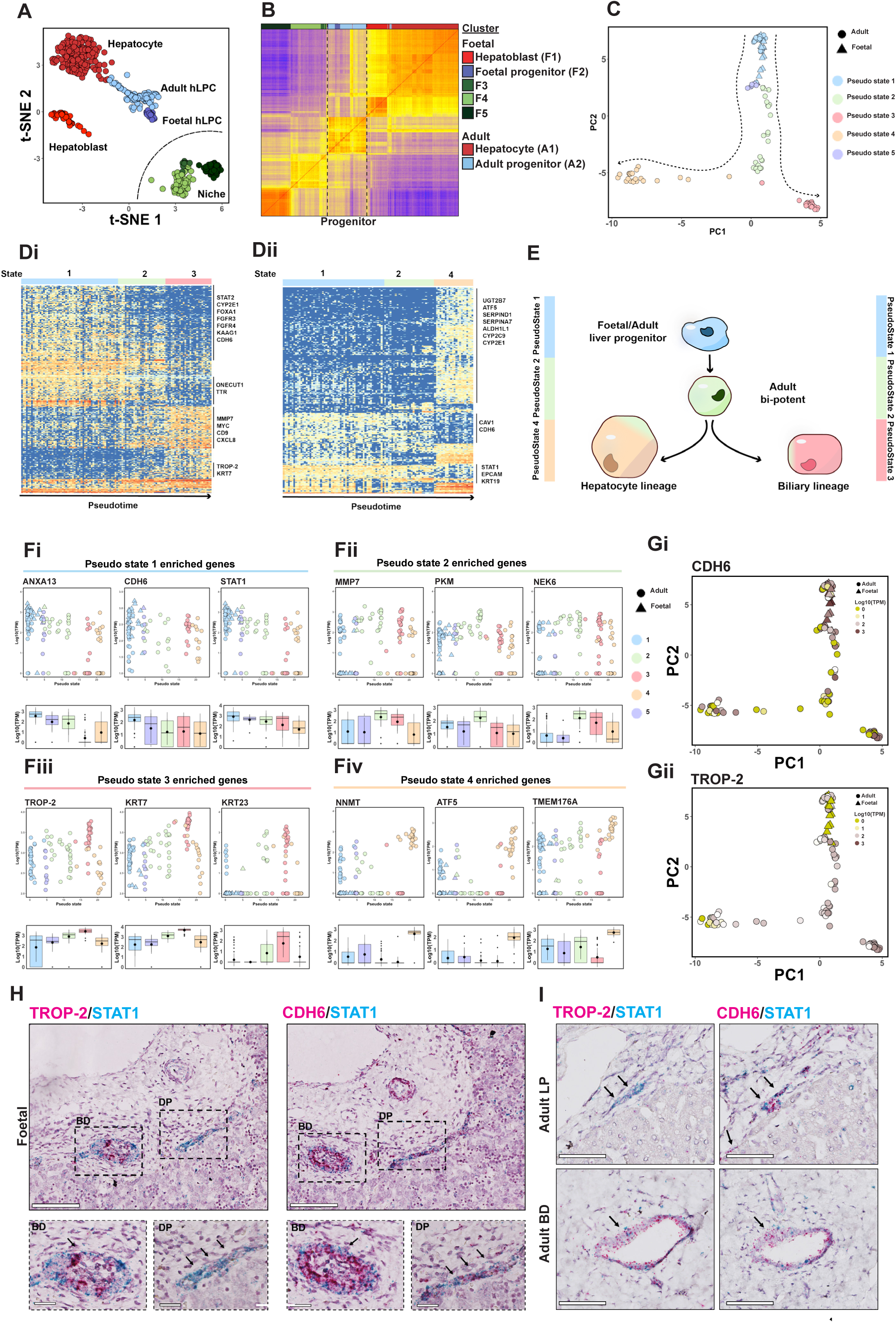
TROP-2 expression differentiates between hLPCs and biliary lineage committed cells. (A) 2D t-SNE representation of all single cells included in this scRNA-seq study. Cells differentially coloured by K-means cluster (adult or foetal). Adult and foetal hLPCs shading (blue) represents single K-means cluster. (B) Heat map showing the spearman correlation coefficient of pairwise comparison between all single cells in data set. The spearman correlation coefficient between samples was calculated based on Log10(TPM) of each gene. Columns are labelled by foetal and adult K-means clusters. (C) PCA plot of Monocle pseudo-time analysis on all cells of foetal and adult hLPC cell clusters, colored by Pseudo state. (D) Heat maps of significantly expressed genes between pseudo states 1,2,3 (i) and 1,2,4 (ii). Gene expression in Log10(TPM). (E) Schematic of hepatic and cholangiocyte liver progenitor lineage based on monocle-derived ‘pseudo state’. (F) Scatter and bar plots of Gene expression Vs Pseudo state for selected genes enriched in pseudo states 1 (i), 2 (ii), 3 (iii) and 4 (iv). (G) PCA plot of Monocle pseudo-time analysis on hLPC cells with overlay of TROP-2 and CDH6 transcript expression. Gene expression in Log10(TPM). (H) Duplex RNA-ISH for TROP-2(red)/STAT1(blue) and CDH6(red)/STAT1(blue) in foetal human bile duct (BD) and ductal plate (DP) structures, detail of the red/blue dots expanded in the square. Scale bars represent 50 μm, 25 μm in blown up squares. (I) Duplex RNA-ISH for TROP-2(red)/STAT1(blue) and CDH6(red)/STAT1(blue) in adult human intra-portal BD and limiting plate (LP) regions. Scale bars represent 25 μm.

### Combined foetal and adult hLPC scRNA-seq analysis identifies TROP-2 as a biliary lineage specification marker

Having confirmed the presence of a CDH6^+^ progenitor population both within foetal and adult human liver tissue, we went on to characterise its potential role in liver development. We first performed 2-D t-SNE distribution and Spearman’s correlation on combined foetal and adult liver scRNA-seq data sets, confirming by K-means clustering that foetal hLPCs are transcriptionally closer to adult hLPCs than any other cell type (Figures 4A-B). Next, to characterise the relationship between foetal and adult hLPCs, we ordered the combined hLPC populations by pseudo-lineage using the R package monocle (Trapnell et al., 2014), to predict the trajectory of hLPC fate. Populations were clustered by ‘pseudo state’ in 2-D space (Figure 4C). Combined hLPCs clustered into five pseudo states. Pseudo state 1 contained all cells of the foetal hLPC population, and a sub-population of adult hLPCs (blue). We observed lineage divergence after pseudo state 2 (green), into states 3 (red) and 4 (yellow). Pseudo state 5 contained a small population of cells that diverged between pseudo state 1 and 2 (Figure 4C). Gene expression analysis revealed significant transcriptional changes from pseudo states 1->2->3 (Figure 4Di), and states 1->2->4 (Figure 4Dii), predicted to represent maturation into cholangiocyte and hepatic lineages, respectively (Figure 4E). Genes enriched in pseudo state 1 included newly defined hLPC markers *CDH6* and *STAT1* (Figure 4 Fi). Pseudo state 3 revealed enrichment of ductal marker genes *KRT7* and *KRT23*, and a novel marker, *TACSDT2* (*TROP-2*) (Figure 4 Fiii). In pseudo state 4, mature hepatic markers *ATF5*, *CYP2C9* and *CYP2E1* were enriched (Figure 4 Fiv). These results suggest that adult hLPCs can be isolated from the liver in three different states, (i) bi-potent, (ii) biliary progenitor or (iii) hepatic progenitor. As expected *CDH6* was significantly enriched in (i) bi-potent cells, whilst *TROP-2* was enriched in (ii) biliary progenitors (Figure 4G). To validate this prediction, we performed duplex RNA-ISH and observed that foetal ductal plate and adult limiting plate regions were negative for *TROP-2* expression, but positive for CDH6 (Figure 4H-I). This suggests *CDH6* positive, *TROP-2* negative cells, define a true bi-potent liver progenitor population, not present in bile ducts.

### Lineage specific markers correlate with hepatocellular carcinoma and intrahepatic cholangiocarcinoma transcriptional signatures

There is no clear consensus on the definition of a ‘cancer stem cell’ or ‘tumour initiating cell’ (Sia et al., 2017). We hypothesized that specific transcriptional changes captured during hepatic and biliary lineage specification, may provide critical insight into the mechanisms of intra-hepatic cancer progression. Utilising a published integrated analysis of differentially expressed genes in hepatocellular carcinoma (HCC) vs intrahepatic cholangiocarcinoma (ICC), we observed a striking correlation with genes enriched during hepatic and biliary lineage specification. 3650 significantly upregulated genes were found in HCC vs ICC, while 3663 genes were significantly downregulated. We compared differentially regulated HCC vs ICC genes to our pseudo lineage bifurcation transcriptional changes (Figure 5 Ai) (Xue et al., 2015). Of the 453 upregulated genes in hepatic progenitors (pseudo state 4) (Figure 5 Aii), 301 were upregulated in HCC, compared to only 9 in ICC (Figure 5 Bi). Of the 201 genes enriched in biliary progenitors (pseudo state 3) (Figure 5 Aii), 90 were upregulated in ICC, compared to only 21 in HCC (Figure 5 Bii) (Supplementary Tables 6-8). These findings demonstrate clearly the close correlation between transcriptional signatures of the biliary and hepatic progenitor state, with respective cancer signatures. We see further evidence of this by comparing biliary and hepatic gene expression with meta-analysis of HCC vs ICC data-sets using the GEPIA web tool (Tang et al., 2017 (Figure 5C-D).

**Figure 5:**
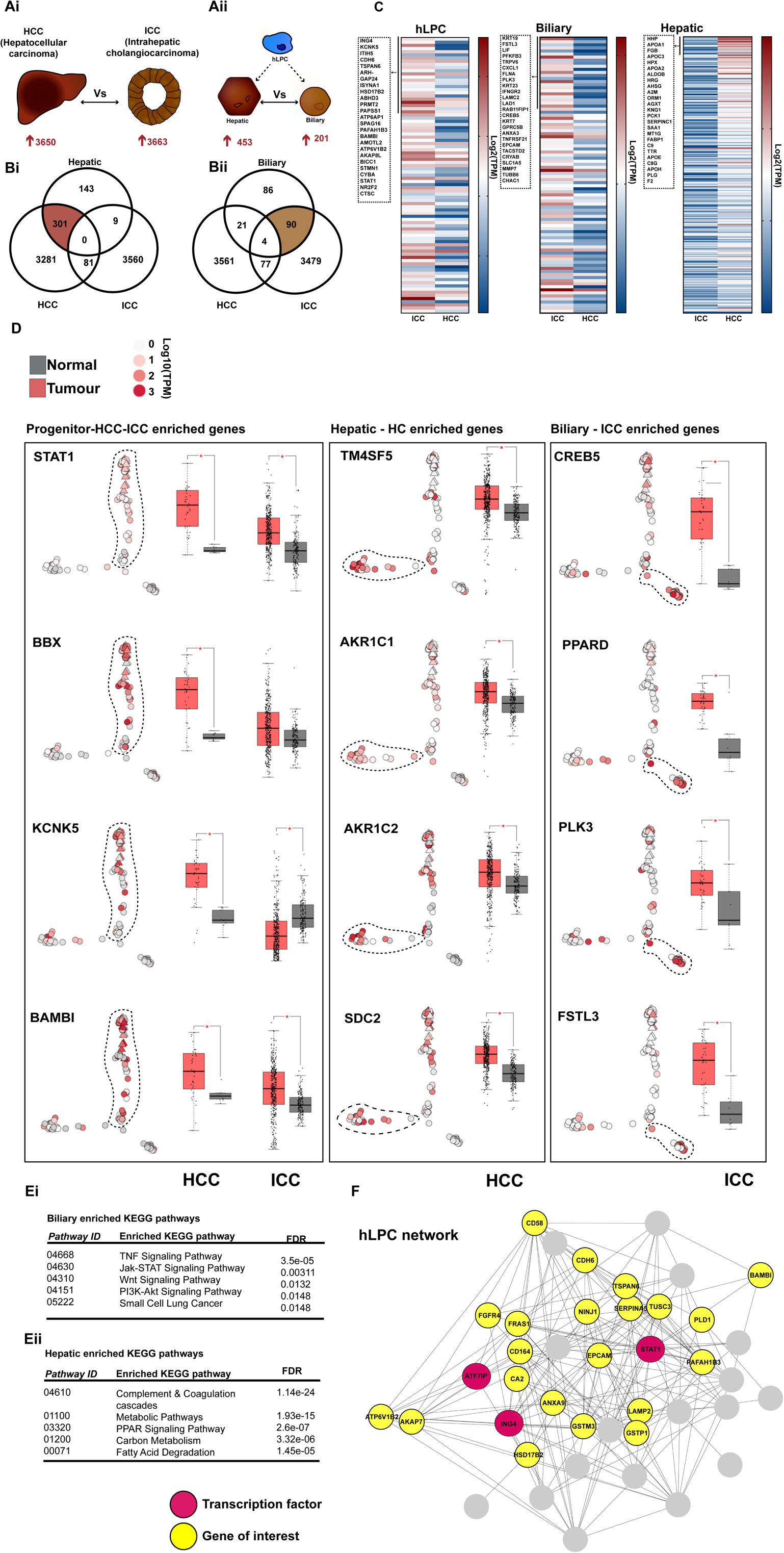
Liver cancer gene signatures correlate with progenitor markers. (A) Schematic of differentially expressed genes (DEGs) between human Hepatocellular Carcinoma (HCC) and Cholangiocarcinoma (ICC) taken from Xue *et al* (Xue et al., 2015)(i), and DEGs between hepatic and biliary pseudo lineage in hLPC scRNA-seq population (ii). (B) Venn diagrams of HCC vs ICC DEGs against pseudo biliary lineage upregulated genes (i) and pseudo hepatic upregulated genes (ii). (C) Heat maps representing the median expression value of DEGs between progenitor, biliary and hepatic gene signatures between GEPIA cholangiocarcinoma (ICC) and hepatocellular carcinoma (HCC) tumour datasets. The names of the top 25 genes in each heat map are shown. Gene expression in log2(TPM). Genes are ranked by expression fold change between progenitor, biliary and hepatic pseudo states. (D) PCA plots of Monocle pseudo-time analysis on all cells of foetal and adult hLPC cell clusters with overlay of selected genes enriched in pseudo state 1 (hLPC), pseudo state 3 (biliary) and pseudo state 4 (hepatic) in Log10(TPM), with box plots of differential gene expression between ICC or HCC samples (red) vs matched normal tissue (grey) in log2(TPM). For ICC comparison: tumour (T) = 36, normal (N) =9. For HCC comparison: T = 369, N = 160. Data acquired from GEPIA interactive web tool (Tang et al., 2017). * represents significant (p-value < 0.01) change in gene expression between tumour and matched tissue samples. (E) KEGG pathways enriched in biliary (i) and hepatic (ii) pseudo lineage correlated with Xue *et al* cancer data sets (Xue et al., 2015). (F) Regulatory network identified by weighted correlation network analysis (WGCNA) in combined foetal and adult hLPC transcriptional profile. Network was constructed on edge weight between nodes. Nodes in red indicate transcription factors in co-expression network, nodes in yellow represents selected genes of interest.

KEGG pathway analysis revealed biliary lineage specification is accompanied by enrichment of TNF, Jak-STAT and WNT signalling (Figure 5 Ei), while hepatic lineage specification is associated with coagulation, metabolic pathways and PPAR signalling (Figure 5 Eii). We performed weighted gene co-expression network analysis (WGCNA) on differentially expressed genes (DEGs) between progenitor, biliary, and hepatic cells, using the R package WGCNA (Figure 5F, Supplementary Figure 5) (Langfelder and Horvath, 2008). As expected, the progenitor co-expression network contained *EpCAM*, *CDH6* and *STAT1*, validating our scRNA-seq predictions of the true hLPC phenotype (Figure 5F). These results demonstrate the close correlation between hLPC fate specification with HCC and ICC gene signatures.

### Identification of an hLPC-specific signature in cancer and chronic liver injury

Having identified a true hLPC population in normal tissue, and observed that lineage specification correlates with gene expression changes in HCC and ICC, we next investigated the distribution of cells with hLPC signatures in cancer and liver injury. In normal adult liver tissue, CDH6 expression is observed in bile ducts and at the limiting plate, whilst TROP-2 is restricted to bile ducts (Figure 6A, Supplementary Figure 6). In HCC tissue with stem-cell like morphological dedifferentiation, *CDH6* expression was observed, whilst *TROP-2* was not. However, cells in ICC were positive for both *CDH6* and *TROP-2* (Figure 6B). Neither CDH6, nor TROP-2, were found in conventional HCC samples. We next looked at *CDH6* and *TROP-2* expression within the context of liver injury and observed acute liver injury to be associated with CDH6^+^/TROP-2^+^ cells (Figure 6C), whilst chronic injury (alcohol) to be associated with CDH6^+^/TROP-2^−^ cells. (Figure 6D). Cumulatively this data suggests that the hLPC population found at the ductal plate, and characterized by CDH6^+^/TROP-2^−^ expression, is involved in dedifferentiated HCC and chronic liver disease. Cholangiocarcinoma and acute liver injury on the other hand, are likely to involve cells from the bile duct as evidenced by observation of CDH6^+^/TROP-2^+^ cells (Figure 7, Supplementary Figure 6). These findings demonstrate that the transcriptional profile of hLPCs identified in our study, have exciting applications in distinguishing between diagnostically challenging liver diseases.

**Figure 6:**
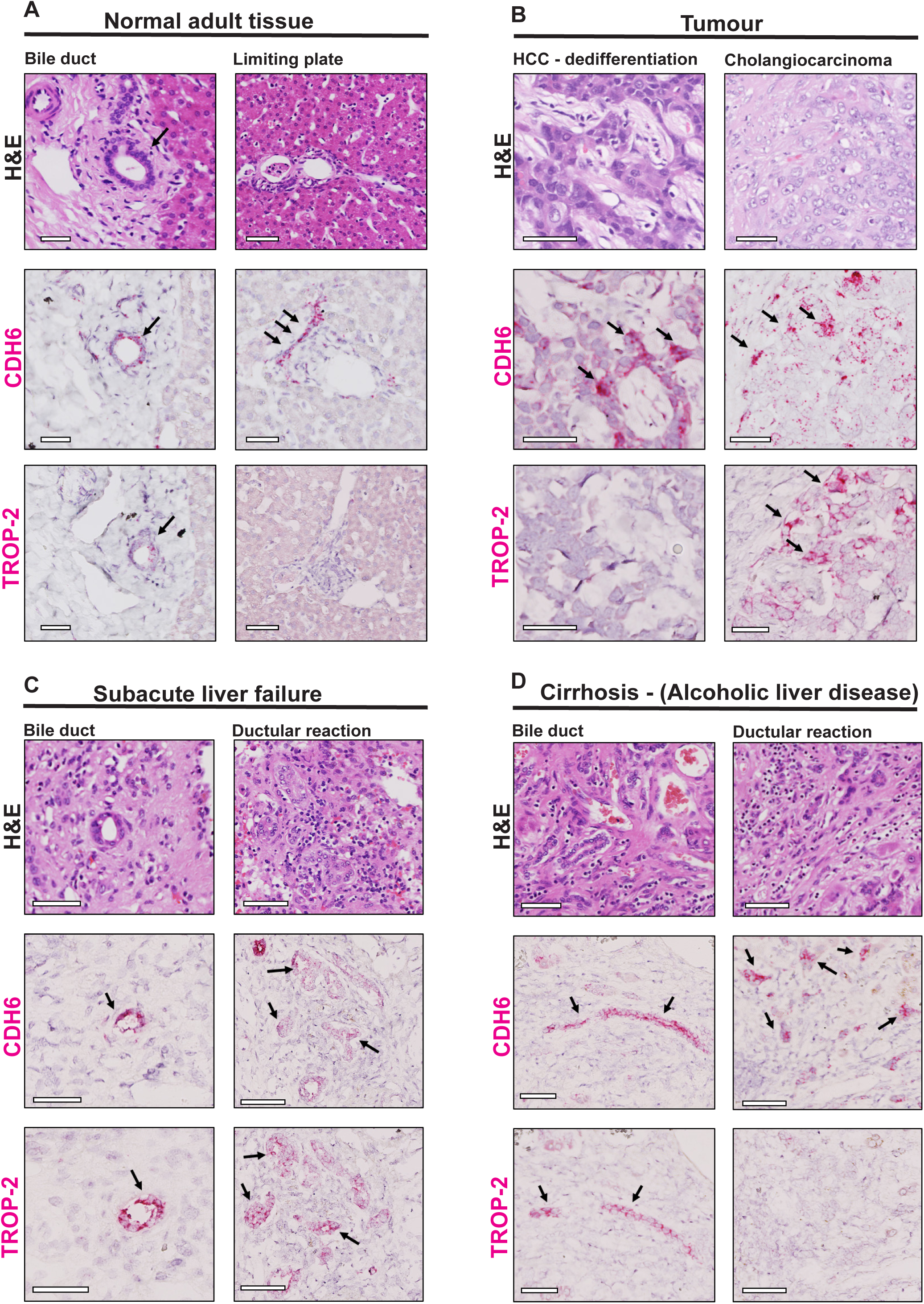
Progenitor subtype signatures correlate with specific human liver injuries. (A) Hematoxylin and eosin (H&E) staining with RNA-ISH for CDH6 and TROP-2 transcript expression in tissue overlay of normal human adult liver tissue. Arrows indicate bile duct and limiting plate structures. (B) H&E staining with RNA-ISH for CDH6 and TROP-2 transcript expression in adult liver tumour samples from hepatocellular carcinoma (HCC) with dedifferentiated morphology and intra-hepatic cholangiocarcinoma (ICC). (C) H&E staining with RNA-ISH for CDH6 and TROP-2 transcript expression in tissue overlay of human adult subacute liver failure bile duct and ductular reactions. (D) H&E staining with RNA-ISH for CDH6 and TROP-2 transcript expression in tissue overlay of human cirrhotic liver bile duct and ductular reactions. All scale bars represent 50 μm.

**Figure 7:**
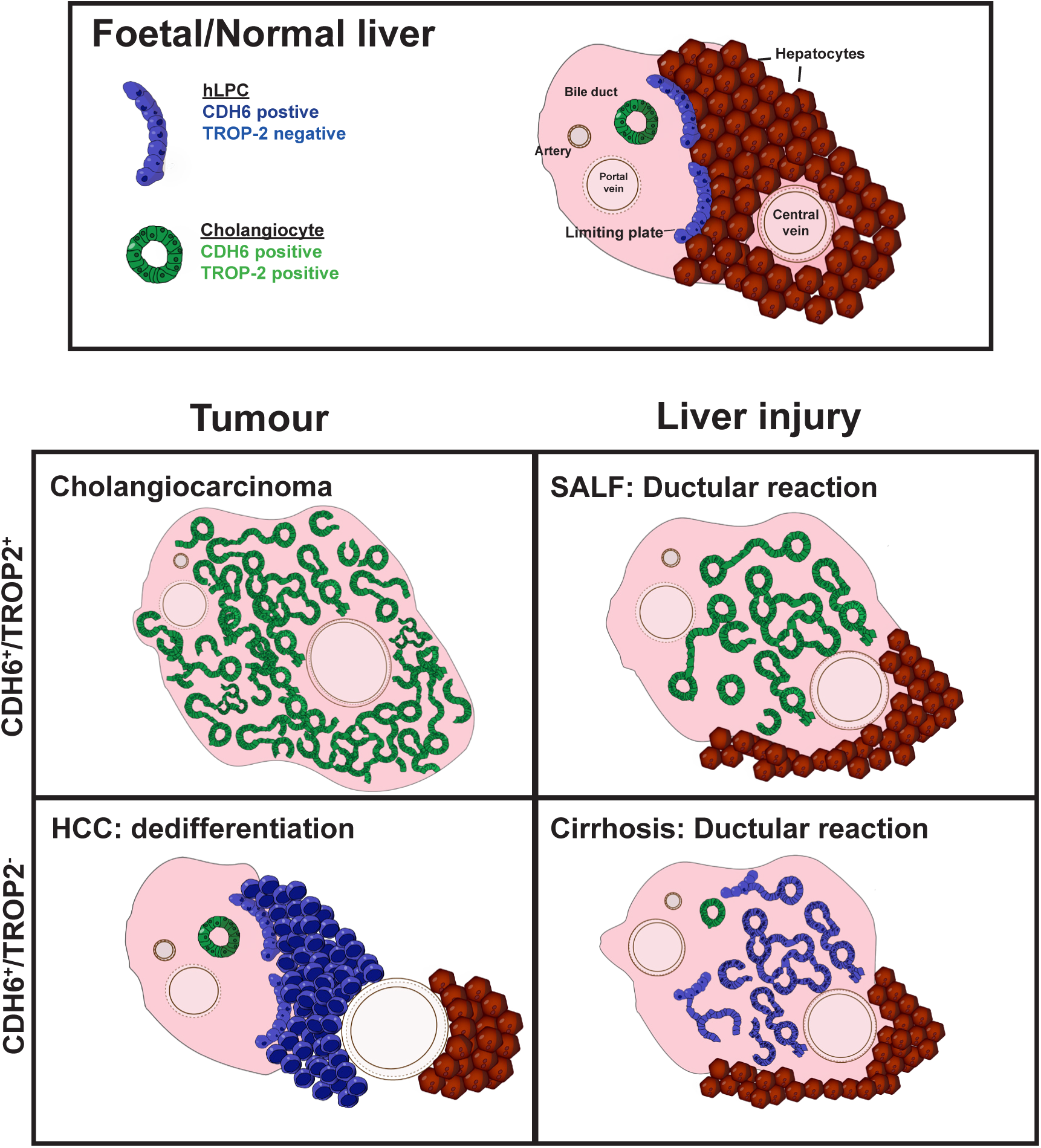
CDH6^+^/TROP-2^−^ cells define a human liver progenitor population normally localized to the limiting plate, present in subtypes of HCC and liver injury. Schematic of human CDH6 and TROP-2 expression in normal, tumour and liver injury models.

### Discussion

In this study, we have captured the molecular identities of three different types of hepatocyte – adult, hepatoblast and progenitor (hLPC) using scRNA-seq. hLPCs localize to the ductal plate of human foetal liver and are retained in its embryological remnant, the limiting plate, of normal adult liver. We identify new surface markers that define this population including CDH6 and FGFR2 along with transcription factors such as STAT1. These cells can be distinguished from cholangiocytes/biliary epithelial cells (BECs) that populate intra-hepatic bile ducts by negative expression of TROP-2. We go on to show that hLPCs may play a putative role within sub-types of chronic liver injury and intra-hepatic cancer.

The current literature has identified putative liver progenitor/stem-cell populations as EpCAM^+^ cells residing in ductal regions or arising from biliary cell dedifferentiation upon injury (Schmelzer et al., 2007, Tarlow et al., 2014, Raven et al., 2017). There is a lack of clarity as to whether the origin and identity of the true hLPC phenotype, compounded by the significant overlap in markers between progenitor/stem-cell populations and cholangiocytes/BECs. Markers including EpCAM, CD24, CD133, CK7, CK19 and SOX9 have been previously associated with both progenitor and BECs in murine and human studies (Yovchev et al., 2007, Mishra et al., 2009, Li et al., 2017). Furthermore, progenitor markers are commonly associated with liver cancer. It is thought these might help in defining the cellular origin of the tumour (progenitor, biliary or mature hepatocyte) and differentiate hepatocellular from cholangiocarcinoma (Suetsugu et al., 2006, Roskams, 2006, Sia et al., 2017). Previous studies lack the single cell resolution necessary to accurately define the origin and identify of the true hLPC phenotype and its functional contribution to development, regeneration in response to injury and misregulation during disease. Our hLPCs, defined at a single cell resolution, on the other hand were positive for conventional markers (*EpCAM*, *CD24*, *CD133*, *KRT7*, *KRT19* and *SOX9*), new markers (*CDH6*, *FGFR2*, *CTNND2*, *STAT1*, *MCAM*, *GPRC5B*) but negative for more mature hepatoblast markers (*AFP*, *CYP3A7* and *DLK1*) (Schwartz et al., 1981, Rowe et al., 2013). These hLPCs were found to reside in the foetal liver ductal plate, and maintained in the adult liver limiting plate. By pseudo-lineage analysis we identified a hLPC bifurcation map which identified TROP-2 as a marker of commitment towards the biliary lineage. We confirmed this using *in situ* RNA hybridization, and showed that a CDH6^+^ hLPC population located at the ductal plate, could be distinguished from cholangiocytes/BECS in bile ducts by the absence of TROP-2 expression.

The power of our experimental approach is highlighted by the finding of hLPC markers CDH6/STAT1 which also appear to be conserved in progenitors across multiple lineages. (Shimazui et al., 2000) (Sagrinati et al., 2006, Kim et al., 2011, Lu et al., 2015, Schmelzer et al., 2007). This supports the idea of a core molecular programme responsible for maintenance of tissue resident stem/progenitor cells (Furuyama et al., 2011, Watt, 2014, Hishikawa et al., 2015). Our data also highlight the important species-specific differences that exist, which likely have led to such confusion in the field. For example, TROP-2 a closely related family member of EpCAM, is thought to be absent in normal tissue and be expressed only at times of injury (Okabe et al., 2009) or cancer (Cubas et al., 2009). Our data clearly shows this is not the case in humans, with TROP-2 being overexpressed in cholangiocarcinoma but also found in bile ducts during normal homeostasis. This raises intriguing questions about the validity of using murine models for human disease research.

Our data provides new knowledge in the direct study of human developmental biology, by molecularly defining the lineage hierarchy from progenitor to adult cell via an intermediate hepatoblast stage (Schmelzer et al., 2007, Schmelzer et al., 2006, Turner et al., 2011). Uncovering the critical regulatory mechanisms driving hepatic maturation will provide an important road map to help unlock the challenges faced in converting pluripotent stem cells to clinically relevant hepatocytes (Baxter et al., 2015). It will also offer new targets for direct cell reprogramming strategies (Nakamori et al., 2017). Finally, our analysis demonstrated remarkable similarity exists between hepatic and biliary progenitors and human HCC and ICC gene expression profiles respectively. This supports the idea that certain forms of cancer may involve reversion to or are driven by cancer stem/progenitor cells and so provides a clinically relevant transcriptional signature to potentially distinguish between diagnostically challenging intra-hepatic tumors.

In summary, we have identified and characterized a true progenitor cell population in human foetal liver, which is retained in normal adult liver and appears to be re-activated in certain types of injury. Further in-depth characterization of the mechanisms regulating the behaviour of this new cell type will afford significant advancement in our understanding of human liver development and disease.

## Methods

### ScRNA-Seq cell sorting and cDNA library preparation

All human tissues were collected with informed consent following ethical and institutional guidelines. Freshly isolated hepatocytes were obtained from Stanford University Hospital. Foetal human livers were obtained from the Human Developmental Biology Resource (HDBR) University College London. Human foetal tissue was dissociated by Collagenase XI enzymatic dissociation for 25 minutes at 37˚C with agitation. Samples were stained with the following primary antibodies, CD235a (349104, FITC, mouse; Biolegend), CD45 (304050, BV711, mouse; Biolegend), EpCAM (324208, APC, mouse; Biolegend), NCAM (362524, PE, mouse; Biolegend), ASGPR1 (563655, PE, mouse, BD Pharmingen™), TFRC (334106, PE mouse, 1:100; Biolegend) all at 1:100 dilution and incubated for 30mins at 4˚C. DAPI (D1306, ThermoFisher Scientific) at 1:1000 dilution was used for live/dead staining. Single-cell sequencing was performed using SmartSeq2 as described (Picelli et al., 2014). Briefly, cells were sorted using a BD FACS Aria II instrument and deposited as single cells into 96-well plates, pre-loaded with lysis buffer (1% Triton X-100, 1mM dNTP, 1μM oligo-dT30, 1:1.2×106 ERCC ExFold RNA spike-in, Recombinant RNase Inhibitor (2313B, Takara Clontech). RNA was converted into complementary (c)DNA using SMARTScribe Reverse Transcriptase (639538, Takara Clontech) and amplified for 21 cycles (Kapa HiFi HotStart ReadyMix 2x, KK2602, KAPA Biosystems). Successful single cell libraries were identified by capillary gel electrophoresis (DNF-474-1000, High Sensitivity NGS Fragment Analysis Kit, AATI) and converted into sequencing libraries using a Nextera XT DNA Sample Preparation Kit (FC-131-1096, Illumina). Barcoded libraries were pooled and subjected to 75 base pair paired-end sequencing on a Illumina NextSeq 2500 instrument. Detail of human foetal and adult liver samples used for sequencing outlined in supplementary table 1.

### DNA sequencing and analysis of Single-Cell Transcriptomes

Raw sequencing reads were aligned using STAR and per gene counts were calculated using HTSEQ (Dobin et al., 2013, Anders et al., 2015). Gene counts were further analyzed using the R package SCATER for pre-processing, quality control and normalization (McCarthy et al., 2017). T-SNE, spearman’s ranks co-efficient, hierarchical clustering, and Student’s t tests were performed using custom scripts in R. T-SNE was performed on log10(TPM) normalized gene counts. User-defined K-means clustering was performed in R. Gene set enrichment analysis (GSEA) was performed on normalized scRNA-seq gene expression data through GSEA software run using the KEGG pathway collection (Subramanian et al., 2005). Pseudo lineage and trajectory analysis was performed in R using the monocle package (Trapnell et al., 2014). Gene co-expression networks were constructed using the blockwiseModules function in the WGCNA package of the R software (Langfelder and Horvath, 2008). Networks were visualized in Cytoscape (Shannon et al., 2003).

### *In Situ* RNA Hybridization (RNA-ISH)

Freshly isolated foetal and adult human liver samples were fixed in 10% formalin buffer saline (HT501128, Sigma Aldrich) for two days then dehydrated and paraffin wax infiltrated using Excelsior™ AS Tissue Processor. After embedding, sections (5µm) were processed for RNA-ISH using the RNAscope 2.5 High Definition (Red, 322350 ACD Bio) according to the manufacturer’s instructions. For single-plex staining the following probes were used; *CDH6 (*Hs-CDH6 403011)*, CD24 (*Hs-CD24 313021)*, FGFR2 (*Hs-FGFR2 311171)*, CTNND2 (*Hs-CTNND2 320269 – custom probe design)*, TACSTD2 (*Hs-TACSTD2 405471) (All ACD Bio). For Duplex RNA-ISH for transcript expression was performed using RNAscope^®^ 2.5 HD Duplex Assay (322435, ACD Bio) according to manufactures instructions using the following custom probes; *STAT1 (*Hs-STAT1 469861, C2 channel change). Slides were counterstained with H&E QS (Vector Laboratories. Inc. Mounted slides were imaged using NanoZoomer (Hamamatsu).

### IF staining

After fixation with 4% (w/v) paraformaldehyde phosphate buffer (PFA) solution (158127, Sigma), cells were blocked and permeabilized in 1% w/v bovine serum albumin (BSA, Sigma-Aldrich), 10% donkey serum (D9663, Sigma Aldrich) and 0.1% Triton (X-100, Sigma Aldrich) for 30 minutes at room temperature (RT). For nuclear antigens, cells were treated with 0.5% Triton (X-100, Sigma Aldrich) for 10 minutes. Primary antibodies, used at the indicated dilutions, CDH6 (AF2715, sheep, 1:50; RnD systems), CK19 (628506, mouse, 1:100; Biolegend), EpCAM (ab7504, mouse, 1:100; abcam) were applied overnight at 4˚C. After washes, cells were incubated with Alexa 647, Alexa 568, Alexa 488 conjugated secondary antibodies (all Life Technologies). Samples were counterstained with NucBlue (R37605, Life Technologies). Images captured with Operetta^®^ (Perkin Elmer).

### Tissue processing and analysis for IF and IHC

For IF staining, foetal liver samples were OCT (23-730-571, ThermoFisher Scientific) embedded, sectioned (5µm) and fixed for 10 minutes in 4% w/v PFA. Primary antibodies were used at the indicated dilutions: CDH6 (AF2715, sheep, 1:50; RnD systems), Claudin-3 (83609S, rabbit, 1:50, Cell Signaling) and STAT1 (610115, mouse, 1:100; Transduction laboratories) at 4˚C overnight. Cells were incubated Alexa 647, Alexa 568, Alexa 488 conjugated secondary antibodies (all Life Technologies) and counterstained with Dapi (D1306, ThermoFisher Scientific). Images were acquired with a Nikon A1 confocal microscope (Nikon Instruments Inc.). Digital images were processed using NIS elements Advanced Research (Nikon) or ImageJ (https://imagej.nih.gov/ij/).

### IHC – Tissue sections

Paraffin-embedded foetal liver tissue were prepared as described for RNA-ISH. After embedding, sections (5µm) were stained using mouse and rabbit specific HRP/AEC (ABC) Detection IHC Kit (abcam) using antibodies against CK19 (ab52625, rabbit, 1:200; abcam), EPCAM (ab7504, mouse, 1:200; abcam), CDH6 (ABIN950438, mouse, 1:100; Antibodies online) and STAT1 (610115, mouse, 1:100; Transduction laboratories) then counterstained with Eosin Y dye (ab146325, abcam). Mounted slides were imaged using a NanoZoomer (Hamamatsu).

### Biliary brush cytology staining

After fixation, cells were blocked and permeabilized in 1% w/v bovine serum albumin (BSA) (A2153, Sigma-Aldrich), 3% donkey serum and 0.1% Triton for 1h at RT. Cells were washed in 0.5% B.S.A, and incubated with the following primary antibodies, CDH6 (AF2715, sheep, 1:50; RnD systems), Claudin-3 (83609S, rabbit, 1:50, Cell Signaling), EPCAM (ab7504, mouse, 1:100; abcam) and KRT7 (ab9021, mouse, 1:100; abcam) overnight at 4˚C. After 3x washes, cells were then incubated with Alexa 647, Alexa 568, Jackson 488 conjugated secondary antibodies (all 1:500, Life Technologies). DAPI was used to label nuclei. Image acquisition was performed using a Nikon A1 inverted confocal microscope (Nikon Instruments Inc.).

### Isolation and expansion of EpCAM^+^/CDH6^+^ cells from human foetal liver

After dissociation, human foetal liver cell suspension was filtered through a 40 µm nylon cell strainer. For FAC sorting, live/dead staining was performed on dissociated cells for 30 minutes at 4˚C in PBS using LIVE/DEAD^®^ Fixable Dead Cell Stain (L10119, Near-red, 1:1000, Thermo scientific). Cells were resuspended in 3% B.S.A with 0.1mM EDTA (15575020, Gibco, Life technologies) and stained for the following conjugated primary antibodies, CD235a (349104, FITC, mouse; Biolegend), CD45 (304050, BV711, mouse; Biolegend), EpCAM (324221, PE/Cy7, mouse, 1:100, Biolegend), Claudin-3 (130-110-834, PE, REA, 1:50, Miltenyl Biotech), CDH6 (130-113-099, APC, REA, 1:50, Miltenyl Biotech) and CD133 (372808, BV 421, mouse, 1:100, Biolegend) for 30 minutes at 4˚C. Fluorescence Minus One (FMO) controls were prepared for each fluorescent channel used. Cells were FAC-sorted on a BD FACS Aria III Fusion and collected in 3% B.S.A with 0.1mM EDTA and 10 µM Y27632 (S1049, Selleckchem), pelleted at 300g for 5 mins and resupended in LE media. FAC sorted cells in LE media were plated directly on to RFP-HUVECS (kindly donated) in endothelial cell growth medium (EGM™ BulletKit™, CC-2517, Lonza) on 0.5µg/ml Matrigel^®^ (354230, Corning) coated plates. RFP-HUVECs were seeded at 15 000 cells per well of a 48 well plate. Co-culture media was maintained at 1:1 ratio of LE: EGM™ BulletKit™ replenished every 24h. Cells were passaged by Trypsin-EDTA (25200056, ThermoFischer Scientific) for 5 minutes at RT and re-plated onto Matrigel^®^.

### F-IHO/F-EHO seeding

Dissociated and filtered foetal liver cells were incubated with anti-CD326 (EpCAM) MicroBeads (130-061-101, Miltenyl Biotech) in 0.5 % BSA with 5U/ml DNASE1 (M0303, N.E.B) and passed through a large cell separation column 2x (130-042-202, miltenyl biotech). Total, EpCAM enriched and EpCAM depleted populations were FACs analyzed for CD235a and EpCAM on a BD Fortessa cell analyzer. EpCAM enriched cells were pelleted at 300 × g for 5 minutes. The cell pellet was resupended in a Matrigel^®^ dome containing ~50,000 cells/dome in one well of a 48-well plate covered in 250μl of LE media. For f-EHO expansion, foetal human gall bladders were obtained from the HDBR network. Tissue samples were washed in Williams E medium (A1217601, Gibco, life technologies) and epithelium scraped off using a surgical blade (adapted from (Sampaziotis et al., 2017)). The supernatant was collected, washed and pelleted at 300 x g. The cell pellet was resuspended in Matrigel^®^ seeded at 1 dome per well of a 48-wellm plate in f-EHO media.

### Liver expansion media

LE media was based on DMEM/F-12, GlutaMAX™ (10565018, Gibco, life technologies) supplemented with 1% N2 supplement (17502048, Gibco, life technologies), 2% B27 supplement (17504044, Gibco, life technologies), 20 mM HEPES (15630080, Gibco, Life technologies), 1.25 mM N-Acetylcysteine (A7250, Sigma), 1% penicillin-streptomycin (15140122, Sigma Aldrich), 1% Insulin:Transferrin: Selenium (ITS) (41400045, Gibco, life technologies), and the growth factors: 50 ng/ml EGF (AF-100-15, Peprotech), 500 ng/ml, RSPO1 (120-38, Peprotech), 100 ng/ml FGF10 (100-26, Peprotech), 25 ng/ml HGF (00-39, Peprotech), 10 mM Nicotinamide (72340, Sigma), 25 µM RepSox (3742, Tocris), 10 µM Forskolin (1099, Tocris), 0.1 μM dexamethasone (1126, R&D Systems), 25 ng/ml Noggin (120-10C, Peprotech), 1:100 Wnt3a (homemade, described in (Willert et al., 2003), and 10 µM Y27632. Hepatic differentiation (HD) media was based on DMEM/F-12, GlutaMAX™ supplement supplemented with 20 mM HEPES, 1.25 mM N-Acetylcysteine (A7250, Sigma), 1% penicillin-streptomycin (15140122, Sigma Aldrich), 1% ITS, 0.1 μM dexamethasone, 10 µM Y27632, 25 ng/ml HGF (00-39, Peprotech), 10 µM DAPT (D5942, Sigma), 2ng/ml Oncostatin-M (295-OM-050, Bio-Techne) and 10ng/ml BMP4 (314-BP-010, Bio-Techne). F-EHOs media was based on DMEM/F-12, GlutaMAX™ supplemented 20 mM HEPES, 1.25 mM N-Acetylcysteine, 1% penicillin-streptomycin, 1% ITS and the growth factors: 50 ng/ml EGF, 500 ng/ml, RSPO1, 0.1 μM dexamethasone, 1:100 WNT3a and 10 µM Y27632. The culture medium was changed every 48 hours.

### Passaging and staining organoids

To split f-IHOs and f-EHOs, Matrigel^®^ was digested with Trypsin-EDTA for 15 minutes at 37 °C. Cell suspension was centrifuged at 300g for 4 minutes, washed once with William’s E medium and resuspended in Matrigel^®^ domes as described above. Organoids were typically passaged at a 1:4 ratio. For IF staining of organoids, Matrigel^®^ was dissolved in cell recovery solution (11543560, ThermoFischer scientific) for 30 minutes at 4˚C and fixed for 30 minutes in 4% w/v PFA. Blocking was performed for 1h in 3% donkey serum, 0.3% Triton and 0.1% DMSO. Primary antibodies were used 4˚C overnight at the indicated dilutions: CDH6 (AF2715, sheep, 1:50; RnD systems), EPCAM (ab7504, mouse, 1:100; abcam), KRT7 (ab9021, mouse, 1:100; abcam), HNF4A (ab92378, rabbit, 1:100; abcam), SOX9 (AF3075, 1:100, R&D Systems), Claudin-3 (83609S, rabbit, 1:50, Cell Signaling), CK19 (ab52625, rabbit, 1:200; abcam), albumin (A80-129A, goat, 1:100; Bethyl). Dapi was used at 1:1000 dilution as a counterstain. Alexa Fluor-conjugated secondary antibodies (all 1:500, Life Technologies). Images were acquired with a Nikon A1 inverted confocal microscope, a Nikon Ti spinning disk confocal microscope (Nikon Instruments Inc.) and a Leica TCS SP8 microscope (Leica Biosystems).

### Isolation of ERCP intra-hepatic biliary brush cytology samples

ERCP samples were collected with informed consent following ethical and institutional guidelines from Denmark Hill Hospital, KCL. Samples were collected in UW^®^ cold storage solution (Belzer), washed 3 x in cold Williams E buffer, and seeded directly into Matrigel^®^ (354230, Corning) domes for 48hs in LE media. For Immunofluorescence, ERCP structures were isolated from Matrigel^®^ in cell recovery solution (354253, Corning™) for 30 minutes at 4˚C, and fixed in 4% w/v PFA. Fixed samples were washed 3X in cold PBS before immunofluorescent staining.

## Acknowledgments

The research was funded/supported by the National Institute for Health Research (NIHR) Clinical Research Facility at Guy’s & St Thomas’ NHS Foundation Trust and NIHR Biomedical Research Centre based at Guy’s and St Thomas’ NHS Foundation Trust and King’s College London. We are grateful to the BRC Flow Cytometry Facility, King’s College Hospital NHS Foundation Trust) for advice and technical assistance. The views expressed are those of the author(s) and not necessarily those of the NHS, the NIHR or the Department of Health. The project has received funding from the European Union’s Horizon 2020 Research and Innovation Programme under the Marie Sklodowska-Curie grant agreement number 705607 We would also like to thank Prof Fiona Watt, Simon Broad and Eamonn Morrison for their support.

## Author Contributions

STR, JS, DW, HN, and SQ designed the experimental approach which was carried out by all other authors, led by JS. Data was analyzed by JS. All authors contributed to the writing of the manuscript which was led by JS.

## References

Aksoy, S., Szumlanski, C. L. & Weinshilboum, R. M. 1994. Human liver nicotinamide N-methyltransferase. cDNA cloning, expression, and biochemical characterization. J Biol Chem, 269, 14835–40.

Anders, S., Pyl, P. T. & Huber, W. 2015. HTSeq−−a Python framework to work with high-throughput sequencing data. Bioinformatics, 31, 166–9.

Ballester, B., Medina-RIVERA, A., Schmidt, D., GonzÀLEZ-PORTA, M., Carlucci, M., Chen, X., Chessman, K., Faure, A. J., Funnell, A. P., Goncalves, A., Kutter, C., Lukk, M., Menon, S., Mclaren, W. M., Stefflova, K., Watt, S., Weirauch, M. T., Crossley, M., Marioni, J. C., Odom, D. T., Flicek, P. & Wilson, M. D. 2014. Multi-species, multi-transcription factor binding highlights conserved control of tissue-specific biological pathways. Elife, 3, e02626.

Baxter, M., Withey, S., Harrison, S., Segeritz, C. P., Zhang, F., Atkinson-DELL, R., Rowe, C., Gerrard, D. T., Sison-YOUNG, R., Jenkins, R., Henry, J., Berry, A. A., Mohamet, L., Best, M., Fenwick, S. W., Malik, H., Kitteringham, N. R., Goldring, C. E., Piper HANLEY, K., Vallier, L. & Hanley, N. A. 2015. Phenotypic and functional analyses show stem cell-derived hepatocyte-like cells better mimic fetal rather than adult hepatocytes. J Hepatol, 62, 581–9.

Cubas, R., Li, M., Chen, C. & Yao, Q. 2009. Trop2: a possible therapeutic target for late stage epithelial carcinomas. Biochim Biophys Acta, 1796, 309–14.

Dabeva, M. D., Petkov, P. M., Sandhu, J., Oren, R., Laconi, E., Hurston, E. & Shafritz, D. A. 2000. Proliferation and differentiation of fetal liver epithelial progenitor cells after transplantation into adult rat liver. Am J Pathol, 156, 2017–31.

Darmanis, S., Sloan, S. A., Croote, D., Mignardi, M., Chernikova, S., Samghababi, P., Zhang, Y., Neff, N., Kowarsky, M., Caneda, C., Li, G., Chang, S. D., Connolly, I. D., Li, Y., Barres, B. A., Gephart, M. H. & Quake, S. R. 2017. Single-Cell RNA-Seq Analysis of Infiltrating Neoplastic Cells at the Migrating Front of Human Glioblastoma. Cell Rep, 21, 1399–1410.

Dobin, A., Davis, C. A., Schlesinger, F., Drenkow, J., Zaleski, C., Jha, S., Batut, P., Chaisson, M. & Gingeras, T. R. 2013. STAR: ultrafast universal RNA-seq aligner. Bioinformatics, 29, 15–21.

Doweiko, J. P. & Nompleggi, D. J. 1991. Role of albumin in human physiology and pathophysiology. JPEN J Parenter Enteral Nutr, 15, 207–11.

Ferrari JÚNIOR, A. P., Lichtenstein, D. R., Slivka, A., Chang, C. & Carr-LOCKE, D. L. 1994. Brush cytology during ERCP for the diagnosis of biliary and pancreatic malignancies. Gastrointest Endosc, 40, 140–5.

Friedman, S. L. 1996. Hepatic stellate cells. Prog Liver Dis, 14, 101–30.

Furuyama, K., Kawaguchi, Y., Akiyama, H., Horiguchi, M., Kodama, S., Kuhara, T., Hosokawa, S., Elbahrawy, A., Soeda, T., Koizumi, M., Masui, T., Kawaguchi, M., Takaori, K., Doi, R., Nishi, E., Kakinoki, R., Deng, J. M., Behringer, R. R., Nakamura, T. & Uemoto, S. 2011. Continuous cell supply from a Sox9-expressing progenitor zone in adult liver, exocrine pancreas and intestine. Nat Genet, 43, 34–41.

Gramignoli, R., Green, M. L., Tahan, V., Dorko, K., Skvorak, K. J., Marongiu, F., Zao, W., Venkataramanan, R., Ellis, E. C., Geller, D., Breite, A. G., Dwulet, F. E., Mccarthy, R. C. & Strom, S. C. 2012. Development and application of purified tissue dissociation enzyme mixtures for human hepatocyte isolation. Cell Transplant, 21, 1245–60.

Hishikawa, K., Takase, O., Yoshikawa, M., Tsujimura, T., Nangaku, M. & Takato, T. 2015. Adult stem-like cells in kidney. World J Stem Cells, 7, 490–4.

Hu, J., Liu, J., Xue, F., Halverson, G., Reid, M., Guo, A., Chen, L., Raza, A., Galili, N., Jaffray, J., Lane, J., Chasis, J. A., Taylor, N., Mohandas, N. & An, X. 2013. Isolation and functional characterization of human erythroblasts at distinct stages: implications for understanding of normal and disordered erythropoiesis in vivo. Blood, 121, 3246–53.

Huch, M., Dorrell, C., Boj, S. F., Van ES, J. H., Li, V. S., Van DE WETERING, M., Sato, T., Hamer, K., Sasaki, N., Finegold, M. J., Haft, A., Vries, R. G., Grompe, M. & Clevers, H. 2013. In vitro expansion of single Lgr5+ liver stem cells induced by Wnt-driven regeneration. Nature, 494, 247–50.

Huch, M., Gehart, H., Van BOXTEL, R., Hamer, K., Blokzijl, F., Verstegen, M. M., Ellis, E., Van WENUM, M., Fuchs, S. A., De LIGT, J., Van DE WETERING, M., Sasaki, N., Boers, S. J., Kemperman, H., De JONGE, J., Ijzermans, J. N., Nieuwenhuis, E. E., Hoekstra, R., Strom, S., Vries, R. R., Van DER LAAN, L. J., Cuppen, E. & Clevers, H. 2015. Long-term culture of genome-stable bipotent stem cells from adult human liver. Cell, 160, 299–312.

Kanamori-KATAYAMA, M., Kaiho, A., Ishizu, Y., Okamura-OHO, Y., Hino, O., Abe, M., Kishimoto, T., Sekihara, H., Nakamura, Y., Suzuki, H., Forrest, A. R. & Hayashizaki, Y. 2011. LRRN4 and UPK3B are markers of primary mesothelial cells. PLoS One, 6, e25391.

Kim, K., Park, B. H., Ihm, H., Kim, K. M., Jeong, J., Chang, J. W. & Cho, Y. M. 2011. Expression of stem cell marker CD133 in fetal and adult human kidneys and pauci-immune crescentic glomerulonephritis. Histol Histopathol, 26, 223–32.

Kuhlmann, W. D. & Peschke, P. 2006. Hepatic progenitor cells, stem cells, and AFP expression in models of liver injury. Int J Exp Pathol, 87, 343–59.

Langfelder, P. & Horvath, S. 2008. WGCNA: an R package for weighted correlation network analysis. BMC Bioinformatics, 9, 559.

Li, B., Dorrell, C., Canaday, P. S., Pelz, C., Haft, A., Finegold, M. & Grompe, M. 2017. Adult Mouse Liver Contains Two Distinct Populations of Cholangiocytes. Stem Cell Reports, 9, 478–489.

Lowes, K. N., Brennan, B. A., Yeoh, G. C. & Olynyk, J. K. 1999. Oval cell numbers in human chronic liver diseases are directly related to disease severity. Am J Pathol, 154, 537–41.

Lu, W. Y., Bird, T. G., Boulter, L., Tsuchiya, A., Cole, A. M., Hay, T., Guest, R. V., Wojtacha, D., Man, T. Y., Mackinnon, A., Ridgway, R. A., Kendall, T., Williams, M. J., Jamieson, T., Raven, A., Hay, D. C., Iredale, J. P., Clarke, A. R., Sansom, O. J. & Forbes, S. J. 2015. Hepatic progenitor cells of biliary origin with liver repopulation capacity. Nat Cell Biol, 17, 971–983.

Mccarthy, D. J., Campbell, K. R., Lun, A. T. & Wills, Q. F. 2017. Scater: pre-processing, quality control, normalization and visualization of single-cell RNA-seq data in R. Bioinformatics, 33, 1179–1186.

Miller, I. J. & Bieker, J. J. 1993. A novel, erythroid cell-specific murine transcription factor that binds to the CACCC element and is related to the Krüppel family of nuclear proteins. Mol Cell Biol, 13, 2776–86.

Mishra, L., Banker, T., Murray, J., Byers, S., Thenappan, A., He, A. R., Shetty, K., Johnson, L. & Reddy, E. P. 2009. Liver stem cells and hepatocellular carcinoma. Hepatology, 49, 318–29.

Nakamori, D., Akamine, H., Takayama, K., Sakurai, F. & Mizuguchi, H. 2017. Direct conversion of human fibroblasts into hepatocyte-like cells by ATF5, PROX1, FOXA2, FOXA3, and HNF4A transduction. Sci Rep, 7, 16675.

Paku, S., Dezso, K., Kopper, L. & Nagy, P. 2005. Immunohistochemical analysis of cytokeratin 7 expression in resting and proliferating biliary structures of rat liver. Hepatology, 42, 863–70.

Petersen, B. E., Zajac, V. F. & Michalopoulos, G. K. 1998. Hepatic oval cell activation in response to injury following chemically induced periportal or pericentral damage in rats. Hepatology, 27, 1030–8.

Philippeos, C., Telerman, S. B., OulÈS, B., Pisco, A. O., Shaw, T. J., Elgueta, R., Lombardi, G., Driskell, R. R., Soldin, M., Lynch, M. D. & Watt, F. M. 2018. Spatial and Single-Cell Transcriptional Profiling Identifies Functionally Distinct Human Dermal Fibroblast Subpopulations. J Invest Dermatol.

Picelli, S., Faridani, O. R., BjÖRKLUND, A. K., Winberg, G., Sagasser, S. & Sandberg, R. 2014. Full-length RNA-seq from single cells using Smart-seq2. Nat Protoc, 9, 171–81.

Proserpio, V., Piccolo, A., Haim-VILMOVSKY, L., Kar, G., LÖNNBERG, T., Svensson, V., Pramanik, J., Natarajan, K. N., Zhai, W., Zhang, X., Donati, G., Kayikci, M., Kotar, J., Mckenzie, A. N., Montandon, R., Billker, O., Woodhouse, S., Cicuta, P., Nicodemi, M. & Teichmann, S. A. 2016. Single-cell analysis of CD4+ T-cell differentiation reveals three major cell states and progressive acceleration of proliferation. Genome Biol, 17, 103.

Rashid, S. T., Humphries, J. D., Byron, A., Dhar, A., Askari, J. A., Selley, J. N., Knight, D., Goldin, R. D., Thursz, M. & Humphries, M. J. 2012. Proteomic analysis of extracellular matrix from the hepatic stellate cell line LX-2 identifies CYR61 and Wnt-5a as novel constituents of fibrotic liver. J Proteome Res, 11, 4052–64.

Raven, A., Lu, W. Y., Man, T. Y., Ferreira-GONZALEZ, S., O’DUIBHIR, E., Dwyer, B. J., Thomson, J. P., Meehan, R. R., Bogorad, R., Koteliansky, V., Kotelevtsev, Y., Ffrench-CONSTANT, C., Boulter, L. & Forbes, S. J. 2017. Cholangiocytes act as facultative liver stem cells during impaired hepatocyte regeneration. Nature, 547, 350–354.

Roskams, T. 2006. Liver stem cells and their implication in hepatocellular and cholangiocarcinoma. Oncogene, 25, 3818–22.

Rowe, C., Gerrard, D. T., Jenkins, R., Berry, A., Durkin, K., Sundstrom, L., Goldring, C. E., Park, B. K., Kitteringham, N. R., Hanley, K. P. & Hanley, N. A. 2013. Proteome-wide analyses of human hepatocytes during differentiation and dedifferentiation. Hepatology, 58, 799–809.

Sagrinati, C., Netti, G. S., Mazzinghi, B., Lazzeri, E., Liotta, F., Frosali, F., Ronconi, E., Meini, C., Gacci, M., Squecco, R., Carini, M., Gesualdo, L., Francini, F., Maggi, E., Annunziato, F., Lasagni, L., Serio, M., Romagnani, S. & Romagnani, P. 2006. Isolation and characterization of multipotent progenitor cells from the Bowman’s capsule of adult human kidneys. J Am Soc Nephrol, 17, 2443–56.

Sampaziotis, F., Justin, A. W., Tysoe, O. C., Sawiak, S., Godfrey, E. M., Upponi, S. S., Gieseck, R. L., De BRITO, M. C., Berntsen, N. L., GÓMEZ-VÁZQUEZ, M. J., Ortmann, D., Yiangou, L., Ross, A., Bargehr, J., Bertero, A., Zonneveld, M. C. F., Pedersen, M. T., Pawlowski, M., Valestrand, L., Madrigal, P., Georgakopoulos, N., Pirmadjid, N., Skeldon, G. M., Casey, J., Shu, W., Materek, P. M., Snijders, K. E., Brown, S. E., Rimland, C. A., Simonic, I., Davies, S. E., Jensen, K. B., Zilbauer, M., Gelson, W. T. H., Alexander, G. J., Sinha, S., Hannan, N. R. F., Wynn, T. A., Karlsen, T. H., Melum, E., Markaki, A. E., Saeb-PARSY, K. & Vallier, L. 2017. Reconstruction of the mouse extrahepatic biliary tree using primary human extrahepatic cholangiocyte organoids. Nat Med, 23, 954–963.

Schaub, J. R., Malato, Y., Gormond, C. & Willenbring, H. 2014. Evidence against a stem cell origin of new hepatocytes in a common mouse model of chronic liver injury. Cell Rep, 8, 933–9.

Schmelzer, E., Wauthier, E. & Reid, L. M. 2006. The phenotypes of pluripotent human hepatic progenitors. Stem Cells, 24, 1852–8.

Schmelzer, E., Zhang, L., Bruce, A., Wauthier, E., Ludlow, J., Yao, H. L., Moss, N., Melhem, A., Mcclelland, R., Turner, W., Kulik, M., Sherwood, S., Tallheden, T., Cheng, N., Furth, M. E. & Reid, L. M. 2007. Human hepatic stem cells from fetal and postnatal donors. J Exp Med, 204, 1973–87.

Schwartz, A. L., Marshak-ROTHSTEIN, A., Rup, D. & Lodish, H. F. 1981. Identification and quantification of the rat hepatocyte asialoglycoprotein receptor. Proc Natl Acad Sci U S A, 78, 3348–52.

Shannon, P., Markiel, A., Ozier, O., Baliga, N. S., Wang, J. T., Ramage, D., Amin, N., Schwikowski, B. & Ideker, T. 2003. Cytoscape: a software environment for integrated models of biomolecular interaction networks. Genome Res, 13, 2498–504.

Shimazui, T., Oosterwijk-WAKKA, J., Akaza, H., Bringuier, P. P., Ruijter, E., Debruyne, F. M., Schalken, J. A. & Oosterwijk, E. 2000. Alterations in expression of cadherin-6 and E-cadherin during kidney development and in renal cell carcinoma. Eur Urol, 38, 331–8.

Sia, D., Villanueva, A., Friedman, S. L. & Llovet, J. M. 2017. Liver Cancer Cell of Origin, Molecular Class, and Effects on Patient Prognosis. Gastroenterology, 152, 745–761.

Subramanian, A., Tamayo, P., Mootha, V. K., Mukherjee, S., Ebert, B. L., Gillette, M. A., Paulovich, A., Pomeroy, S. L., Golub, T. R., Lander, E. S. & Mesirov, J. P. 2005. Gene set enrichment analysis: a knowledge-based approach for interpreting genome-wide expression profiles. Proc Natl Acad Sci U S A, 102, 15545–50.

Suetsugu, A., Nagaki, M., Aoki, H., Motohashi, T., Kunisada, T. & Moriwaki, H. 2006. Characterization of CD133+ hepatocellular carcinoma cells as cancer stem/progenitor cells. Biochem Biophys Res Commun, 351, 820–4.

Tanaka, M., Okabe, M., Suzuki, K., Kamiya, Y., Tsukahara, Y., Saito, S. & Miyajima, A. 2009. Mouse hepatoblasts at distinct developmental stages are characterized by expression of EpCAM and DLK1: drastic change of EpCAM expression during liver development. Mech Dev, 126, 665–76.

Tang, Z., Li, C., Kang, B., Gao, G. & Zhang, Z. 2017. GEPIA: a web server for cancer and normal gene expression profiling and interactive analyses. Nucleic Acids Res, 45, W98–W102.

Tarlow, B. D., Pelz, C., Naugler, W. E., Wakefield, L., Wilson, E. M., Finegold, M. J. & Grompe, M. 2014. Bipotential adult liver progenitors are derived from chronically injured mature hepatocytes. Cell Stem Cell, 15, 605–18.

Terada, T. 2013. Ductal plate in hepatoblasts in human fetal livers: I. ductal plate-like structures with cytokeratins 7 and 19 are occasionally seen within human fetal hepatoblasts. Int J Clin Exp Pathol, 6, 889–96.

Trapnell, C., Cacchiarelli, D., Grimsby, J., Pokharel, P., Li, S., Morse, M., Lennon, N. J., Livak, K. J., Mikkelsen, T. S. & Rinn, J. L. 2014. The dynamics and regulators of cell fate decisions are revealed by pseudotemporal ordering of single cells. Nat Biotechnol, 32, 381–386.

Treutlein, B., Brownfield, D. G., Wu, A. R., Neff, N. F., Mantalas, G. L., Espinoza, F. H., Desai, T. J., Krasnow, M. A. & Quake, S. R. 2014. Reconstructing lineage hierarchies of the distal lung epithelium using single-cell RNA-seq. Nature, 509, 371–5.

Turner, R., Lozoya, O., Wang, Y., Cardinale, V., Gaudio, E., Alpini, G., Mendel, G., Wauthier, E., Barbier, C., Alvaro, D. & Reid, L. M. 2011. Human hepatic stem cell and maturational liver lineage biology. Hepatology, 53, 1035–45.

Wang, B., Zhao, L., Fish, M., Logan, C. Y. & Nusse, R. 2015. Self-renewing diploid Axin2(+) cells fuel homeostatic renewal of the liver. Nature, 524, 180–5.

Watt, F. M. 2014. Mammalian skin cell biology: at the interface between laboratory and clinic. Science, 346, 937–40.

Willert, K., Brown, J. D., Danenberg, E., Duncan, A. W., Weissman, I. L., Reya, T., Yates, J. R. & Nusse, R. 2003. Wnt proteins are lipid-modified and can act as stem cell growth factors. Nature, 423, 448–52.

Xue, T. C., Zhang, B. H., Ye, S. L. & Ren, Z. G. 2015. Differentially expressed gene profiles of intrahepatic cholangiocarcinoma, hepatocellular carcinoma, and combined hepatocellular-cholangiocarcinoma by integrated microarray analysis. Tumour Biol, 36, 5891–9.

Yanger, K., Knigin, D., Zong, Y., Maggs, L., Gu, G., Akiyama, H., Pikarsky, E. & Stanger, B. Z. 2014. Adult hepatocytes are generated by self-duplication rather than stem cell differentiation. Cell Stem Cell, 15, 340–349.

Yovchev, M. I., Grozdanov, P. N., Joseph, B., Gupta, S. & Dabeva, M. D. 2007. Novel hepatic progenitor cell surface markers in the adult rat liver. Hepatology, 45, 139–49.

Yovchev, M. I., Grozdanov, P. N., Zhou, H., Racherla, H., Guha, C. & Dabeva, M. D. 2008. Identification of adult hepatic progenitor cells capable of repopulating injured rat liver. Hepatology, 47, 636–47.

